# Gene encoding CC-NBS-LRR protein on rye chromosome 1RS confers wheat stripe rust resistance

**DOI:** 10.1101/2024.10.04.616747

**Authors:** Chunhui Wang, Yanan Chang, Mian Wang, Jing Wang, Chang Liu, Chaolan Fan, Congyang Yi, Chen Zhou, Jing Yuan, Wuyun Yang, Dengcai Liu, Tao Wang, Yang Liu, Xingguo Ye, Fangpu Han

## Abstract

Stripe rust, a globally widespread disease, stands as one of the most significant threats to wheat cultivation. The 1BL/1RS translocation, renowned for its robust resistance to both rust and powdery mildew, has historically played an important role in wheat breeding and production. The gene for resistance to stripe rust on the 1RS is known as *Yr9* and plays an important role in the production of wheat, but over the course of long-term breeding had lost its resistance due to the evolution of stripe rust towards greater and greater virulence. In this paper, we cloned the stripe rust resistance gene, *Yr9*, from triticale by genetic mapping approach. The *Yr9* encodes a typical nucleotide-binding leucine-rich repeat (NLR) protein. Both transgenic and overexpression of *Yr9* in highly stripe rust susceptible wheat varieties conferred complete resistance to the stripe rust races CYR17 and partial resistance to the stripe rust races CYR32, CYR33, and CYR34. In addition, the *Yr9* allele in the 1BL/1RS translocation line also showed the same level of resistance to stripe rust. Both two alleles loses resistance when deployed in the field or inoculated with mixed physiological races collected from the field. Our findings provide valuable insights for breeders to strategically incorporate disease resistance genes and provides a foundation for further understanding how pathogenic bacteria might evolve to evade recognition via NLR type proteins.

**Significance:** The 1BL/1RS translocation between wheat and rye is the most successful case of exogenous gene application in plant genetic improvement and has been used in wheat breeding for over 50 years. Here we report the cloning of a stripe rust resistance gene *Yr9* located on rye chromosome 1RS using a triticale population. The *Yr9* encodes a coiled-coil nucleotide-binding site leucine-rich repeat (CC-NBS-LRR) protein that show complete resistance to the stripe rust races CYR17 and partial resistance to the stripe rust races CYR32, CYR33, and CYR34, albeit demonstrating susceptibility under field conditions. Our findings position *Yr9* as an ideal candidate gene to study the mechanism of inactivation of disease resistance genes as a result of pathogen evolution.

## Introduction

Wheat is one of the most important food crops in classic and modern agriculture, feeding about one-third of the world’s population, and providing about 20 percent of the protein and calories among all human foods (1, 2). Stripe rust is a widespread foliar wheat disease caused by *Puccinia striiformis* f. sp. *tritici* (*Pst*) in many countries and regions, and its prevalence severely threatens global food supplies via both crop destruction (yield losses by 10% to 20%) as well as significant reductions in wheat quality (3–6). As a major wheat producer in the world, in most years, China’s wheat yield is greatly harmed by by stripe rust (7). Therefore, identifying ideal genes and applying them towards new wheat varieties with stripe rust resistance is the most economically sensible and environmentally friendly way to manage this disease.

In the past two or three centuries, wheat was directionally domesticated according to humanity requirements for available traits and high yield, which resulted in the loss of many potentially useful genes and correspondingly a significant reduction of genetic diversity (8, 9). To our knowledge, most wild or semi-wild wheat relatives such as rye (*Secale cereale* L.), *Aegilops* spp., *Haynaldia villosa*, *Thinopyrum* spp., *Agropyron cristatum*, *Leymus mollis*, generally possessed high genetic diversity and good environmental adaptability, which can greatly contributed to the genetic improvement of wheat (10–15). Among them, rye is one of the most valuable genus for wheat breeding. In 1950s, the 1RS chromosome arm from the German rye ‘Petkus’ carrying many available resistance genes to a wide range of diseases such as *Pm8*, *Sr31*, *Lr26*, and *Yr9* was introduced into wheat by distant hybridization (16, 17). The 1RS chromosome arm not only benefited yield but also contributed to increasing environmental adaptability of wheat (18). In 1971, some Russian and Eastern European wheat cultivars (like Kavkaz, Aurora, Lovrin 10, and Predgornia 2) containing 1RS·1BL translocation chromosome were introduced into China, and in time became widely used in wheat breeding (19). By 1990s, 70% of wheat cultivars released in China belonged to 1RS·1BL translocation lines (20, 21). However, the over-reliance on *Yr9* for stripe rust resistance led to the generation of new virulent *Pst* races in China under high evolutionary pressure, and this gene can no longer endow wheat with any resistance to these new *Pst* races (22).

Triticale (×*Triticosecale* Wittmack) is a synthetic plant species containing the genomes of both rye and durum wheat, generated by interspecific hybridization between the two species and chromosome doubling. The plant is much less heterozygous than rye because of triticale undergoing chromosome doubling and self-pollinating, which is a unique advantage over rye for gene mapping using triticale populations.

In this paper, a triticale F_2_ population from the cross of a resistant line and a susceptible line was used to map stripe rust resistance gene *Yr9* by disease resistance phenotyping, Bulked segregant RNA sequencing (BSR-seq) analysis, genome resequencing, and marker development.

## Results

### Genetic identification of stripe rust resistance genes on chromosome 1RS in triticale line L20191212

The two triticale lines L20191212 and 4100 used in this study differed considerably on stripe rust resistance (Fig. 1A). Line L20191212 was near-immune to stripe rust, with obvious hypersensitive flecks on leaves but no uredia developed. On the other hand, line 4100 was susceptible to stripe rust, with few hypersensitive flecks but uredia all over the leaves. Genomic *in situ* hybridization (GISH) assays clearly showed that the two materials contained 7 pairs of rye chromosomes and 14 pairs of wheat chromosomes, and no obvious chromosomal aberrations were found (Fig. 1B). Stripe rust assays in the 224 F_2_ plants between the two lines showed that 173 plants were nearly immune and 51 plants were susceptible to the disease (Fig. S1A). The inheritance behavior of the resistance from line L20191212 was followed 3:1 ratio (χ2_3:1_ = 0.595, P > 0.2), suggesting that the stripe rust resistance in L20191212 is controlled by a single dominant gene (Fig. S1B).

**Figure 1.**
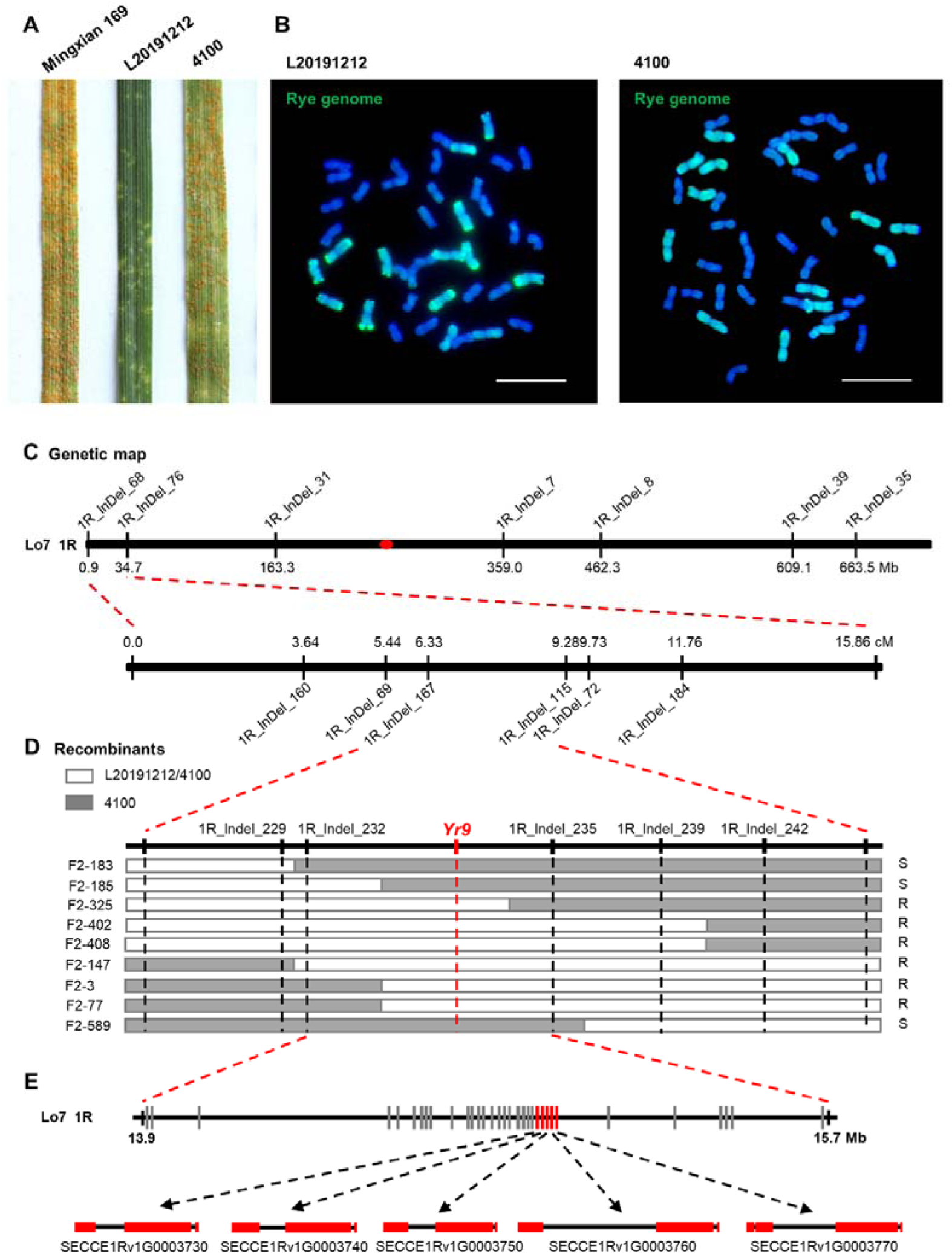
The gene(s) for stripe rust resistance in hexaploid triticale L20191212 is located on chromosome 1RS. (A) Evaluation of stripe rust resistance in triticale L20191212 and 4100. Common wheat cultivar Mingxian 169 was used as a susceptible control. (B) L20191212 and 4100 karyotype identified by GISH. The genomic DNA of *Secale cereale* (green) were used as probe. Chromosomes were stained with DAPI (blue). Scale bar = 10 μm. (C) Genetic map of *Yr9* based on 224 F_2_ plants from cross L20191212 × 4100. The distribution of InDel markers on chromosome 1R and their physical locations based on the Lo7 reference genome. (D) Recombinant haplotypes with L20191212 chromosome segments determined by genotyping. White rectangles indicate heterozygous chromosome regions of L20191212/4100; Gray rectangles represent 4100 chromosomes; R. Resistant; S. susceptible. (E) Physical map of *Yr9*. Red boxes represent NLR genes and grey boxes represent non-NLR genes. The second row shows its gene structure, with red rectangles representing exons.

BSR-seq analysis indicated that the stripe rust resistance gene is likely to be located on chromosome 1R, which was confirmed by using the molecular markers located on 1R (Fig. S2). To further map this gene, we developed a series of InDel (insertion and deletion) markers located at different positions on 1R using the genome resequencing data of L20191212 and 4100 (Dataset S1). Consequently, 13 InDel markers were identified to be located on 1R, and a linkage map was constructed (Fig. 1C). Of these, two markers of 1R_Indel_167 and 1R_Indel_115 were most closely linked to the stripe rust resistance gene in L20191212. For the fine mapping of *Yr9*, we genotyped additional 684 F_2_ plants using five InDel markers, screened nine recombinant haplotypes, and pinpointing *Yr9* between markers 1R_Indel_232 and 1R_Indel_235, which corresponded to 1R: 13.9-15.7 Mb of the Lo7 reference genome (Fig. 1D).

### Screening of candidate genes for *Yr9*

Within the aforementioned 1.8 Mb genome region, we identified 33 high confidence genes, with nine of them expressed (Fig. 1E, Dataset S2). Based on the expression profiles of these genes over a range of times after stripe rust inoculation (Fig. 2A), five proximate genes - SECCE1Rv1G0003730, SECCE1Rv1G0003740, SECCE1Rv1G0003750, SECCE1Rv1G0003760 and SECCE1Rv1G0003770 – clearly showed a significantly higher expression level in L20191212 than in 4100. We then analyzed the expression differences of the five genes in the resistant and susceptible groupings of the F_2_ population, and found that the expression levels of SECCE1Rv1G0003760 and SECCE1Rv1G0003770 were very high in the resistant group but very low (almost no expression) in the susceptible group (Fig. 2A). Additionally, after stripe rust infection, the expression of SECCE1Rv1G0003760 was relatively stable, with a slight decrease during 0-12 h, then increased in the following 60 h while the expression of SECCE1Rv1G0003770 was continuously increased during the first 24 h, and significantly decreased after 48 h (Fig. 2B). Both SECCE1Rv1G0003760 and SECCE1Rv1G0003770 encode typical CNL type proteins, so we characterized the two genes as candidate genes for *Yr9* (Fig. 2C).

**Figure 2.**
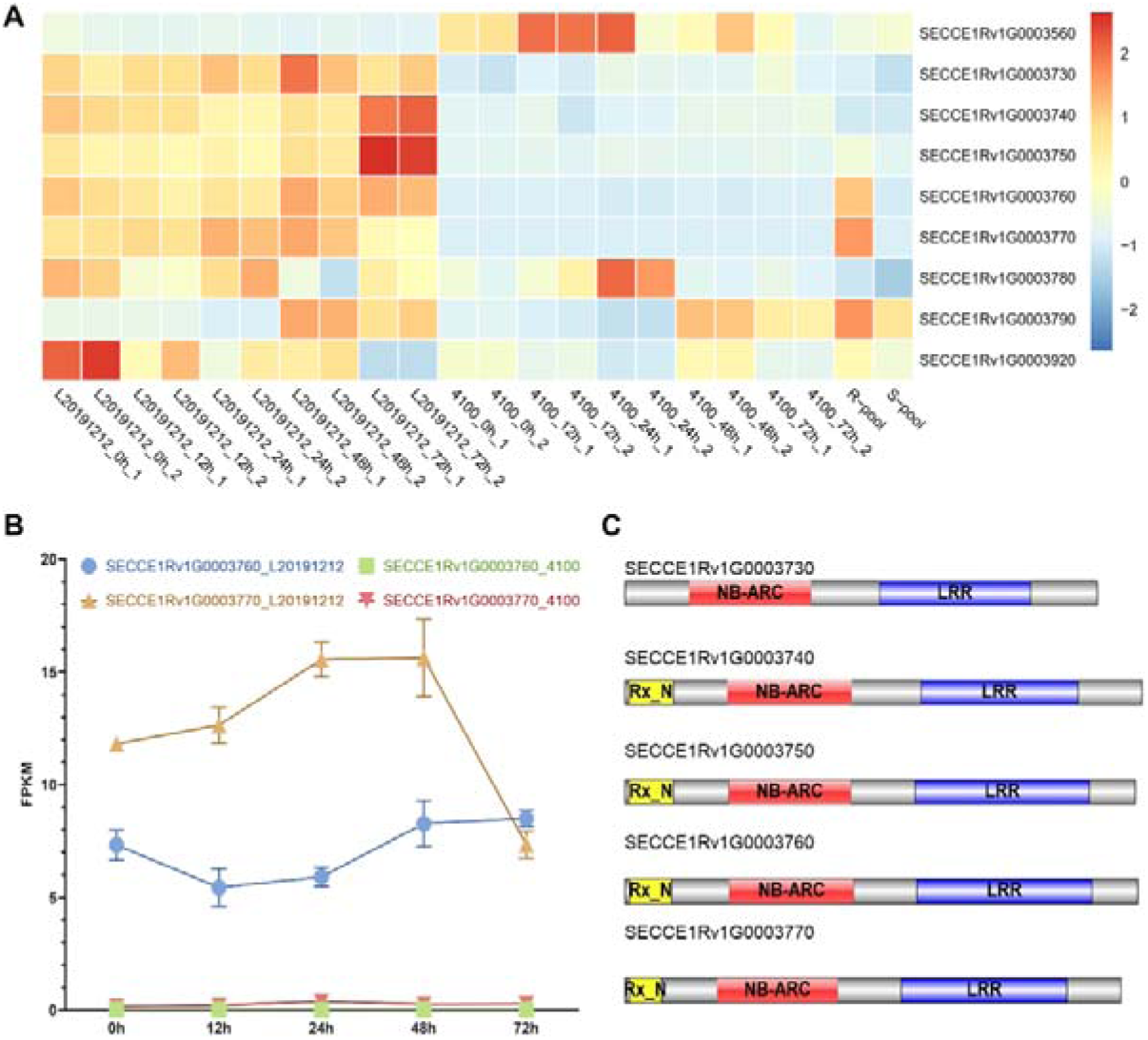
Screening for *Yr9* candidate genes. (A) The heatmap shows the expression of all expressible genes in the interval. The samples were collected at five timepoints (0, 12, 24, 48 and 72 h) after inoculation with CYR32, CYR33 and CYR34 for L20191212 and 4100, respectively, with two replicates for each sample. R-pool: Resistance pool in BSR-seq analysis; S-pool: Susceptibility pool in BSR-seq analysis. (B) Line graph of candidate gene expression under stripe rust induction in resistant and susceptible parents. (C) Predicted domains of all NLR proteins in the *Yr9* interval. CC: coiled-coil, NBS: nucleotide-binding, LRR: leucine-rich repeat.

### Candidate gene SECCE1Rv1G0003760 being functionally validated as *Yr9*

To verify the function of the two candidate genes, we overexpressed them in two wheat cultivars, Fielder and Yannong 19, which are highly susceptible to stripe rust (Fig. 3A). We tested the T_1_ transgenic seedlings with the two candidate genes for stripe rust resistance, and found that all the transgenic plants containing *SECCE1Rv1G0003770* were highly susceptible to stripe rust (Fig. 3B). Meanwhile, the transgenic plants containing *SECCE1Rv1G0003760* showed near-immune resistance to stripe rust races CYR17, with a few hypersensitive flecks without any uredia on the leaves, demonstrating that SECCE1Rv1G0003760 is *Yr9* (Fig. 3B). To rule out any potential unforeseen effects of the ubiquitin promoter on stripe rust resistance, we generated more transgenic plants of *Yr9* using its native promoter in the wheat cultivar Fielder and found that the transgenic plants driven by the *Yr9* native promoter showed the same stripe rust resistance to the transgenic plants regulated by the ubiquitin promoter (Fig. 4 B and C). Remarkably, *Yr9* exhibited partial resistance to the prevailing physiological *Pst* races in China (CYR32, CYR33, and CYR34), ranging from moderately resistant to near-immune (Fig. S3). However, it’s worth noting that *Yr9* lost resistance to the mixed stripe rust races collected in the field that was virulent to *Yr9*, indicating that *Yr9* does not have broad-spectrum resistance (Fig. S4). Similarly, in the Sichuan field, the transgenic positive plants of *Yr9* showed high susceptibility to stripe rust (Fig. S5).

**Figure 3.**
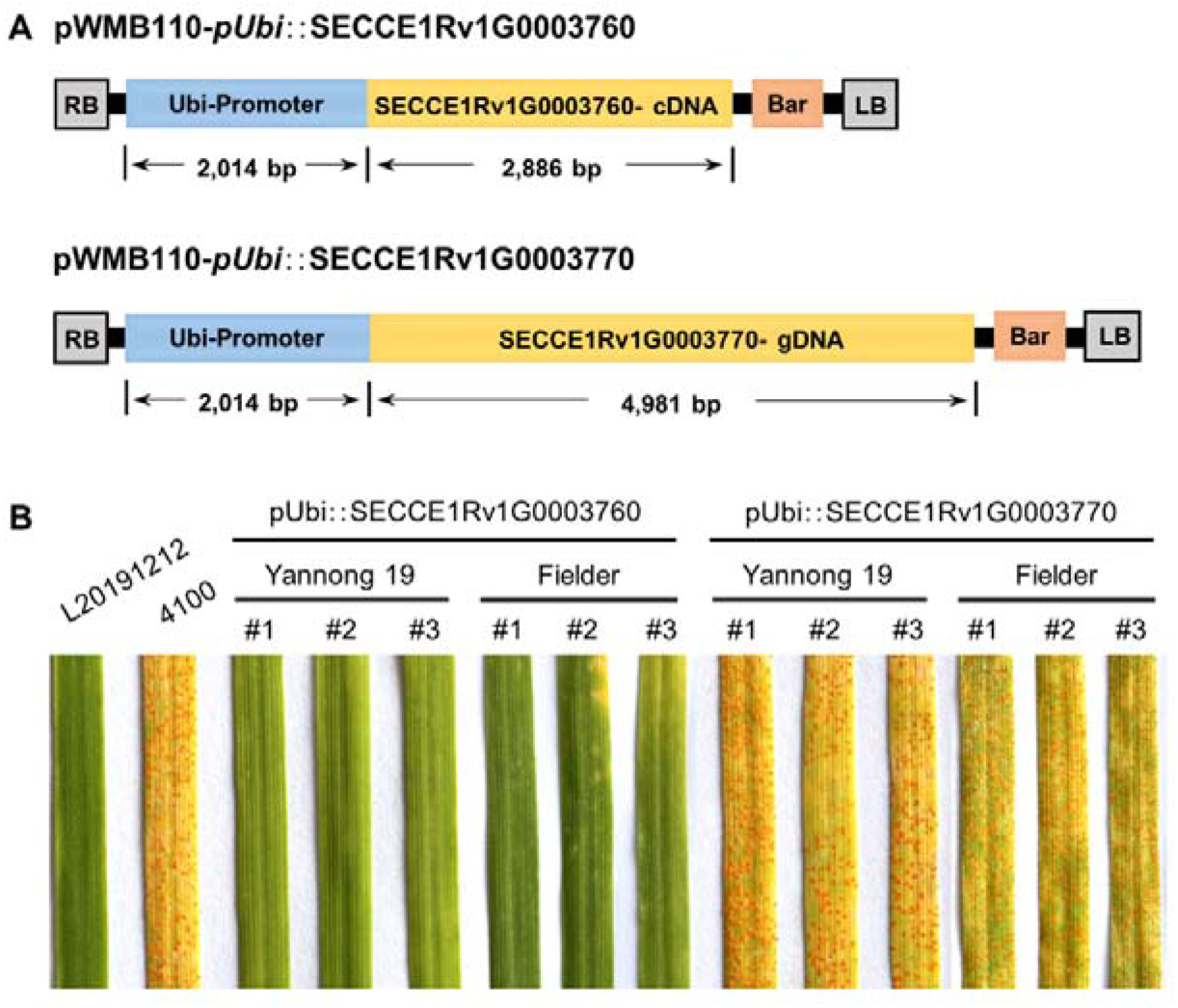
Functional validation of *Yr9*. (A) The CDS of *SECCE1Rv1G0003760* (2,886 bp) and the genomic sequence of *SECCE1Rv1G0003770* (4,981 bp) used for transformation. (B) Infection types of T_1_ generation plants of *SECCE1Rv1G0003760* and *SECCE1Rv1G0003770* in response to stripe rust race CYR17.

**Figure 4.**
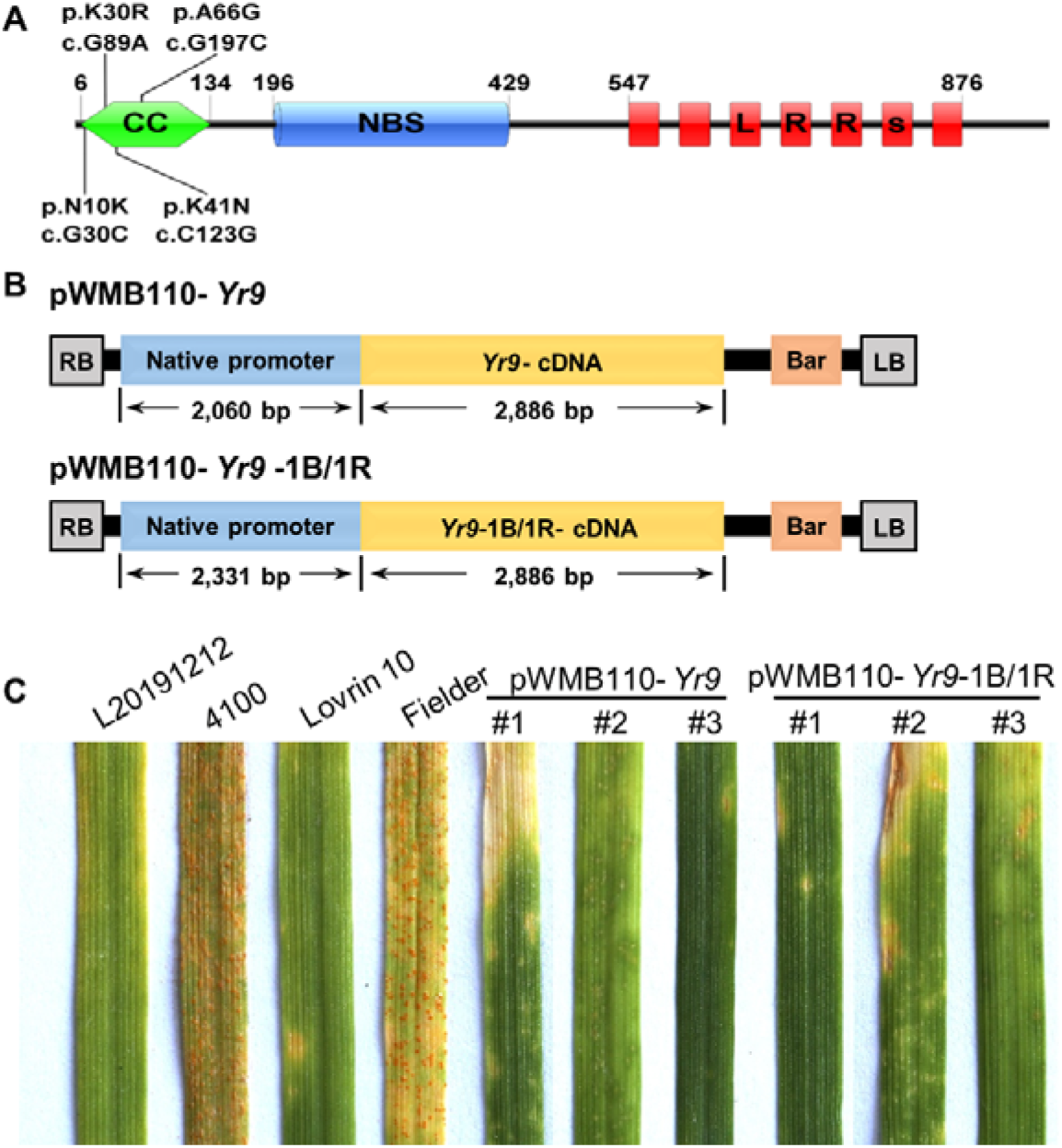
*Yr9* allele in the 1BL/1RS translocation line resistance to stripe rust. (A) Differential sites of *Yr9* with *Yr9*-1B1R. (B) The CDS of *Yr9* (2,886 bp) and presumed promoter (2,060 bp) used for transformation. The CDS of *Yr9*-1B1R (2,886 bp) and presumed promoter (2,331 bp) used for transformation. (C) Performance of the *Yr9* and *Yr9*-1B1R for resistance to stripe rust. The stripe rust race used in this experiment was stripe rust race CYR17.

### *Yr9* alleles in the 1BL/1RS translocation lines having similar functions of stripe rust resistance

We cloned the full-length sequences of *Yr9* alleles in 34 1BL/1RS translocation lines and found that the basic sequences were almost the same in all the cultivars tested. Additionally, 19 SNPs were identified among the allele of *Yr9* in the 1BL/1RS translocation cultivars in comparison to the *Yr9*, ten in intron I and nine in exon I, resulting in four amino acid changes (Fig. 4A, Fig.S6, and Fig.S7). We tentatively named the *Yr9* allele in the 1BL/1RS translocation as *Yr9*-1B/1R. To verify whether the *Yr9*-1B/1R has the same function of stripe rust resistance, we also introduced it into Fielder (Fig. 4B). The T_1_ transgenic plants showed the same stripe rust resistance to the transgenic plants harboring *Yr9* after being inoculated with the stripe rust races, suggesting that the four amino acid differences of *Yr9*-1B/1R do not affect its function on stripe rust resistance (Fig. 4C, Fig. S3). Then, we further cloned the full-length sequences of *Yr9* in 21 triticale, and found that 19 of them were consistent with the sequence in the Lo7 reference genome and only one was consistent with the sequence of *Yr9*-1B/1R, suggesting that the *Yr9* used in wheat production in the past decades worldwide was a relatively rare type from rye (Fig. S7).

## Discussion

Long-term domestication via selection for high yield has resulted in low genetic diversity in wheat, leading to its weak resistance to many pests and diseases. To improve wheat resistance to various diseases, geneticists and breeders have made great efforts to introduce chromosomes or segments carrying available resistance genes into wheat from its closed genera through distant hybridization (23, 24). Using this strategy, which is really a feat rarely implemented in other plant species, has greatly enriched wheat genetic diversity. An example of the most successful case of distant hybridization is the Rye chromosome 1RS, which endows wheat with both resistance to rust and mildew and with yield improvement (25, 26).

Translocation and introgression lines are the main vectors for the application of exogenous genes in wheat breeding and production, but the cloning of exogenous genes in these materials is always challenging. The main reason for that issue is due to the recombination limitation between the alien chromosome and its corresponding wheat chromosomes. Recently developed technologies based on mutant sequencing, such as MutRenSeq, MutChromSeq, and MutRNASeq, have shown great advantages and potential applications for cloning exogenous disease resistance genes (27–29). However, the acquisition of a large number of mutants is time-consuming and labor-intensive. Many previous attempts have been made to clone genes from rye using triticale populations, but with little success (30–32). Our work demonstrates that it is feasible to use triticale populations to clone from rye and provides a viable method for exogenous gene cloning.

*Yr9* was resistant to almost all stripe rust races at all stages when it was introduced into China in the 1970s (33, 34). However, the massive cultivation of 1BL/1RS translocation cultivars greatly strengthened the evolutionary pressure on *Pst*, and virulent races soon emerged (35). According to the latest results published, *Yr9* has lost resistance to many of the *Pst* physiological races in the field, but *Yr9* is resistant to two of the most common stripe rust races (CYR32 and CYR34) in China (22), which is consistent with our current results in this study. Interestingly, the pyramiding combinations of *Yr9* and other genes (e.g., *Yr17*, *Yr30*) have been reported to enhance wheat resistance to stripe rust (36). Therefore, although the use of *Yr9* alone is not desirable in wheat production, the combination application of a few stripe rust resistance genes with *Yr9* is still valuable.

Nowadays, the 1BL/1RS translocation varieties are becoming less valuable in wheat breeding and grain production due to the reduced resistances to stripe rust and powdery mildew as well as the negative impact on flour quality (37, 38). But, the potential of 1BL/1RS translocation still can be revitalized if the following issues could be addressed in the future: 1) identifying and modifying key amino acids in *Yr9* and *Pm8* and improve their disease resistance by using synthetic biology and gene editing techniques; 2) to precisely introduce new disease resistance genes onto the 1BL/1RS chromosome by using large fragment insertion techniques; 3) to knock out the secalin genes on the 1BL/1RS chromosome for improving flour quality. We plan to address these queries and hope that this work would inspire others to similarly journey forth.

## Materials and methods

### Plant materials

Hexaploid triticale lines L20191212 and 4100 were kindly provided by, respectively, Prof. Xingfeng Li at Shandong Agricultural University and Prof. Zhongfu Ni at China Agricultural University. The hybrids of L20191212 and 4100 were self-crossed one generation to form a F_2_ population. The highly susceptible stripe rust wheat cultivar Minxian 169 kept in our group was used to culture the *Pst* races. Two highly susceptible stripe rust cultivars, Fielder and Yannong 19, were maintained in our laboratory. These were used as transformation recipients to generate transgenic plants. Detailed information of the triticale lines and the 1BL/1RS translocation cultivars used in this study are listed in Dataset S3.

### Stripe rust assays

Seeding-stage stripe rust assays were carried out at the Stripe Rust Reaction Platform at the Institute of Genetics and Developmental Biology, Chinese Academy of Sciences. Specific methods of stripe rust inoculation were performed as described in a previous article (23). The stripe rust races used in this investigation included CYR17, CYR32, CYR33, CYR34, and Field mixed *Pst* races (collected at Qingcheng Experimental Station of Gansu Academy of Agricultural Sciences, Tianshui City, Gansu, China).

### Cytogenetic assays

For Genome in situ hybridization (GISH) experiment, fresh root tips grown for 2 d were collected and treated with nitrous oxide for 2-3 h, followed by 90% acetic acid for 8-10 min to fix chromosomes. Metaphase chromosome preparations and probe labelling were performed according to a previously published protocol (39). In this study, the rye Lo7 genome was labelled with Alexa Fluor-488-5-dUTP by nick translation. Microscopy and visualization were conducted as a described operation (40).

### BSR-Seq, RNA-seq and bioinformatics analysis

Thirty extremely resistant and 30 extremely susceptible plants in the L20191212/4100 F_2_ population were sampled to construct a resistant bulk and a susceptible bulk, respectively. At five timepoints (0, 12, 24, 48, and 72 h) after stripe rust inoculation with a mixture of three races CYR32, CYR33 and CYR34, the leaves of L20191212 and 4100 were collected and total RNA was extracted as described in the miRNeasy Mini Kit (Cat: 217004; TIANGEN). Consequently, the total RNA was sequenced on the Illumina NovaSeq 6000 platform (ANOROAD GENE TECHNOLOGY) to generate 125 bp pairs of reads. All raw data was uploaded to the National Genomics Data Center under accession number CRA019067. Due to the lack of a reference genome for hexaploid triticale, we combined the *Secale cereale* Lo7 Reference Genome v1.0 (41) and the *Triticum turgidum* Svevo reference genome v1.0 (42) as the reference genome for triticale for the following analyses. Trimmomatic-0.36 was used to remove low quality and adapter sequences from raw reads (43).

For BSR-seq, reads were aligned to the triticale reference genome to generate Bam files using STAR v2.7.9a (44). Subsequently, the MarkDuplicates and AddOrReplaceReadGroups modules of the picard v2.5.0 software (http://broadinstitute.github.io/picard/) were used to remove duplicates generated from PCR amplification and sort bams. SplitNCigarReads module of the GATK v4.0.11.0 software was then used to split reads at intronic regions (45). Calling SNPs used the HaplotypeCaller module of GATK v3.2-2 and filtering and plotting used the R package QTLseqr (46).

Transcriptome analyses were performed according to previously published protocols (47). Briefly, data from triticale parents L20191212 and 4100 after different times of stripe rust inoculation were individually aligned to the triticale genome using hisat2-2.0.4 (48). Transcripts were assembled and gene expression levels predicted using StringTie (49), the following differential gene expression analysis using the R package Ballgown (50). The R package pheatmap was used to plot expression heatmaps for genes with FPKM>1 in at least one sample.

### InDel marker development

Fresh leaves of two hexaploid triticale parents, L20191212 and 4100, were used to extract DNA using the CTAB method. The DNA was sequenced using the Illumina NovaSeq 6000 platform (ANOROAD GENE TECHNOLOGY) with 150 Gb of data per sample (accession number CRA019068). Reads alignment and genomic variation calling were performed as previously published paper with some adjustments (51). Cleaned data processed by Trimmomatic-0.36 was aligned to the triticale reference genome using BWA-MEM (52). The parameters for filtering the obtained InDels are set as follows: “QD < 10.0, FS > 200.0, QUAL < 100.0” and “ReadPosRankSum < −20.0”. The filtered InDels greater than or equal to 5 bp were selected for primer design using the NCBI Primer Design Tool (https://www.ncbi.nlm.nih.gov/tools/primer-blast/), and *Tricum aestivum* (taxid:4565) was selected in Exclusion Organism option. Polymorphic primers that are co-dominant between L20191212 and 4100 can be used for genetic mapping. A total of 247 primer pairs were designed in this paper, of which 37 pairs met the requirements and 18 pairs were selected for genetic mapping.

### Genetic mapping of *Yr9*

Genetic mapping of *Yr9* was performed using an F2 population consisting of 684 plants from a cross between L20191212 and 4100. We first genotyped 224 F2 plants, including 173 resistant and 51 susceptible plants, using 13 co-dominant markers to construct a linkage map. The genetic map of *Yr9* was constructed using QTL IciMapping Version 4.2 (53). The chi-squared test (χ2) was used to determine the goodness of fit of the observed segregation rate to the theoretical Mendelian rate. To fine mapping of *Yr9*, 460 plants were genotyped using five markers (1R_Indel_229, 1R_Indel_232, 1R_Indel_235, 1R_Indel_239 and 1R_Indel_242) to screen for key recombinant haplotypes.

### Protein domain prediction

The genome sequences and protein sequences of related genes were retrieved from the rye Lo7 reference genome in the Ensembl plants database (54), and the domains of the proteins were predicted online via the interpro website (http://www.ebi.ac.uk/interpro/).

### Plasmid Constructs and Wheat transformation

The CDS of SECCE1Rv1G0003760 (2886 bp) and the genomic DNA of SECCE1Rv1G0003770 (4981 bp) including the entire coding region and introns were amplified in triticale L20191212 using KOD OneTM PCR Master Mix (Toyobo, Japan). These two DNA fragments were ligated behind ubiquitin (*Ubi*) promoter in vector *pWMB110* using the EasyGeno Assembly Cloning Kit (TianGen, Beijing, China). The two plasmids were introduced into stripe rust susceptible common wheat cultivars Fielder and Yannong 19 via *Agrobacterium tumefaciens* (strain GV3101) mediated transformation. A total of 4946 bp of SECCE1Rv1G0003760 native promoter (2060 bp) and gene coding region (2886 bp) were ligated into the vector *pWMB110* with removed the ubiquitin promoter. In addition, the homologous sequences of SECCE1Rv1G0003760 in the 1RS·1BL translocation Aikang 58 including the native promoter (2331 bp) and the gene coding region (2886 bp) were also ligated into the vector *pWMB110*. Both of these plasmids were also introduced into the wheat cultivars Fielder by an Agrobacterium-mediated transformation. All primers used in the amplification sequences and construction vector in this paper are listed in Dataset S2.

## Acknowledgement

This work was supported by the National Natural Science Foundation of China (NSFC32494473012 and 31991212) and the “Biological Breeding-National Science and Technology Major Project” (2023ZD04025).

## Supporting Information includes

**Figure S1.**
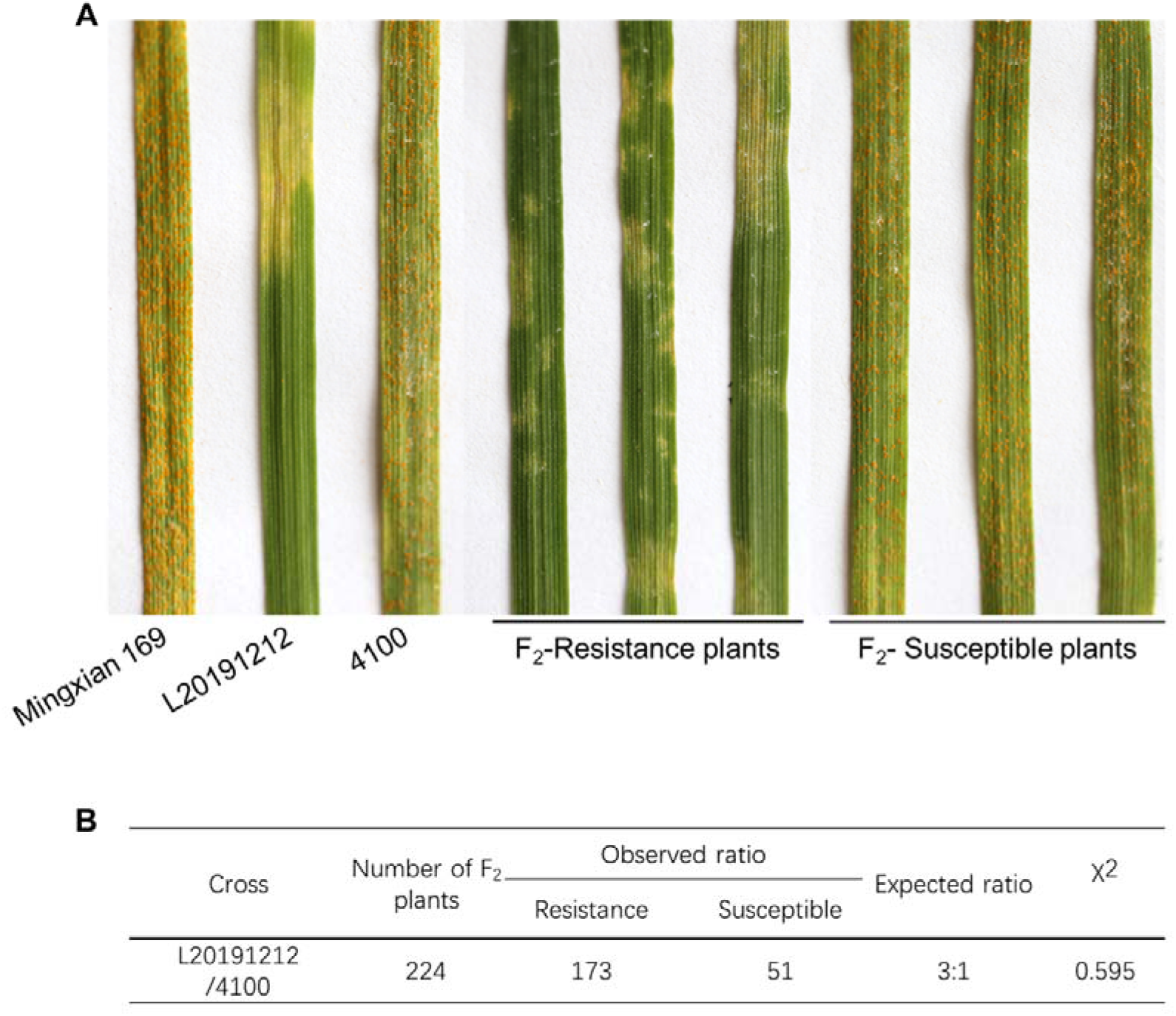
Stripe rust resistance in triticale L20191212 F_2_ population. (A) Stripe rust resistance in representative plants of L20191212 F_2_ population and parents. (B) Statistical results of stripe rust resistance in the F_2_ population.

**Figure S2.**
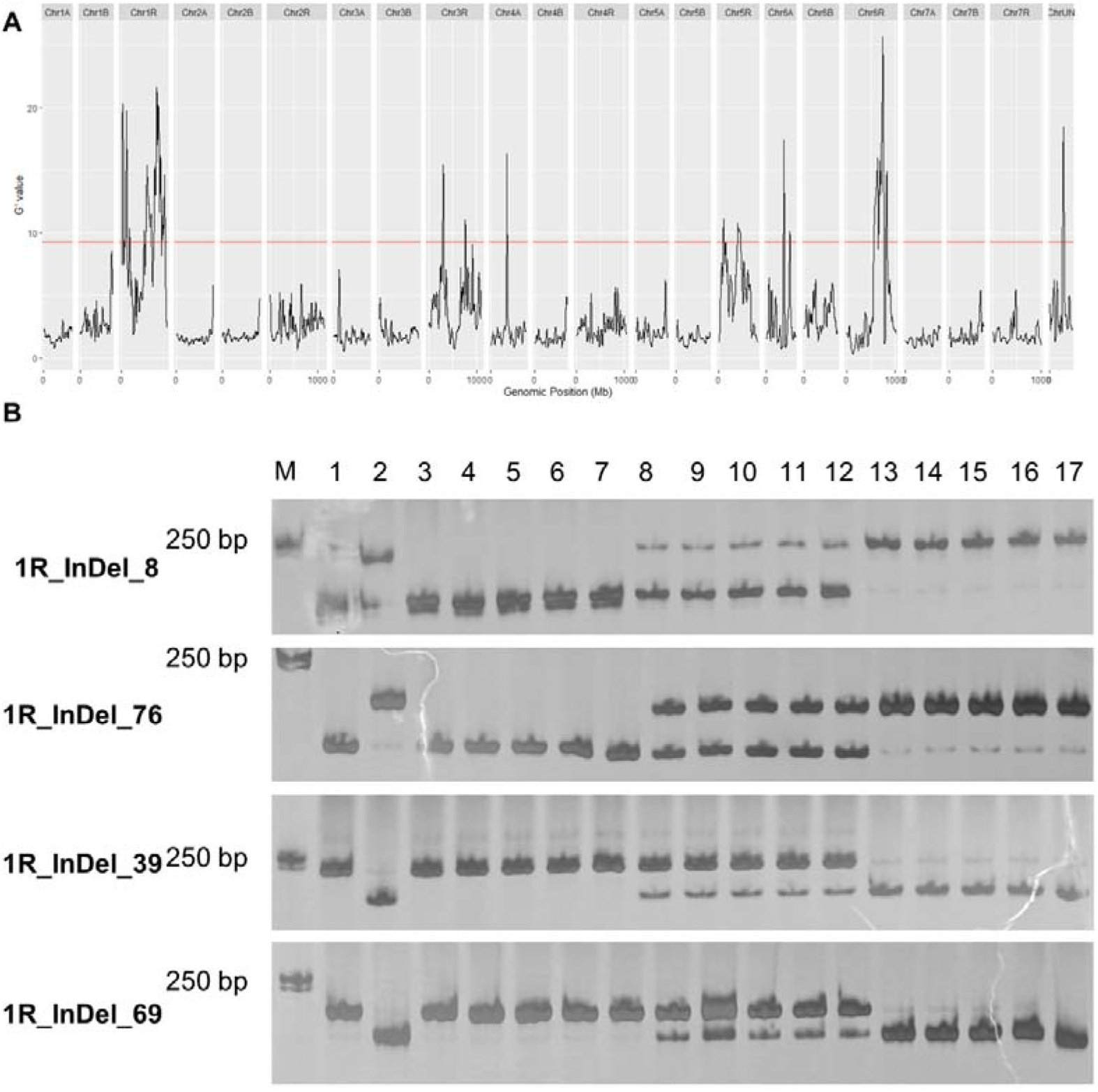
1R chromosome of L20191212 resistance to stripe rust. (A) BSR analysis of L20191212 using the G’value method. (B) Genotyping of the F2 population using 1R InDel markers. M: Marker, 1:L20191212, 2:4100, 3-7: the homozygous of F_2_ plants, 8-12: the heterozygous of F_2_ plants, 13-17: susceptible plants.

**Figure S3.**
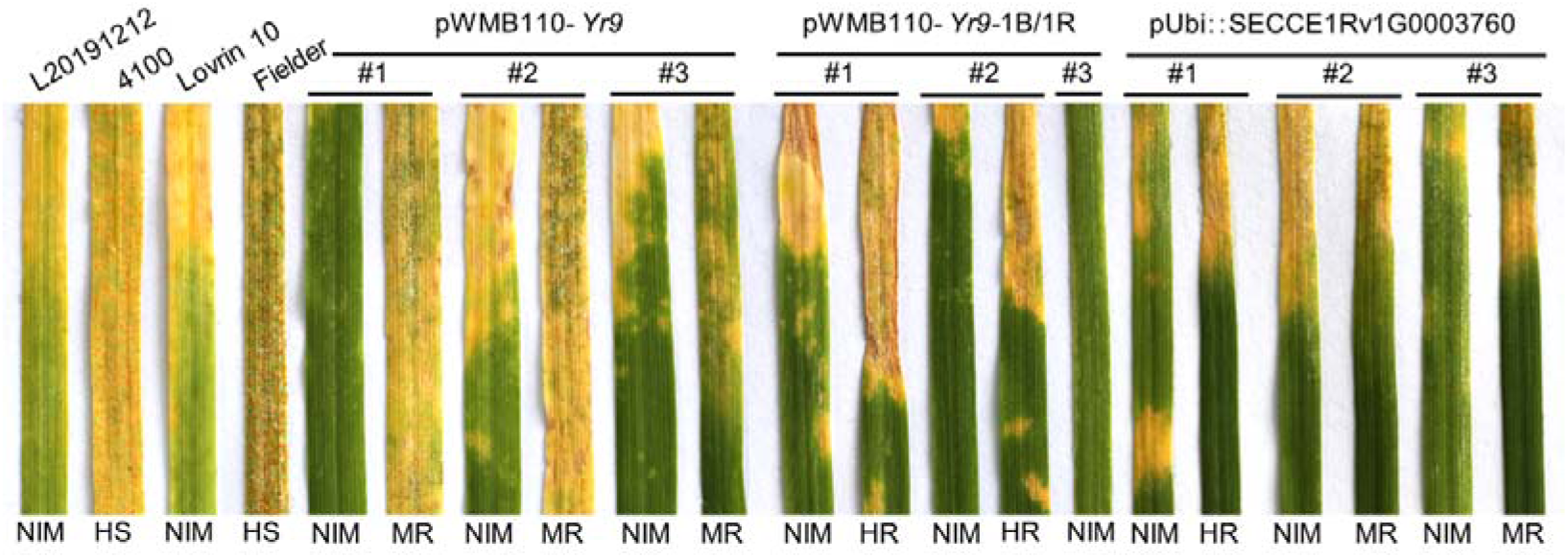
Resistance of *Yr9* to stripe rust races CYR32, CYR33 and CYR34. Representative phenotypes are shown, and resistance levels are labelled below.

**Figure S4.**
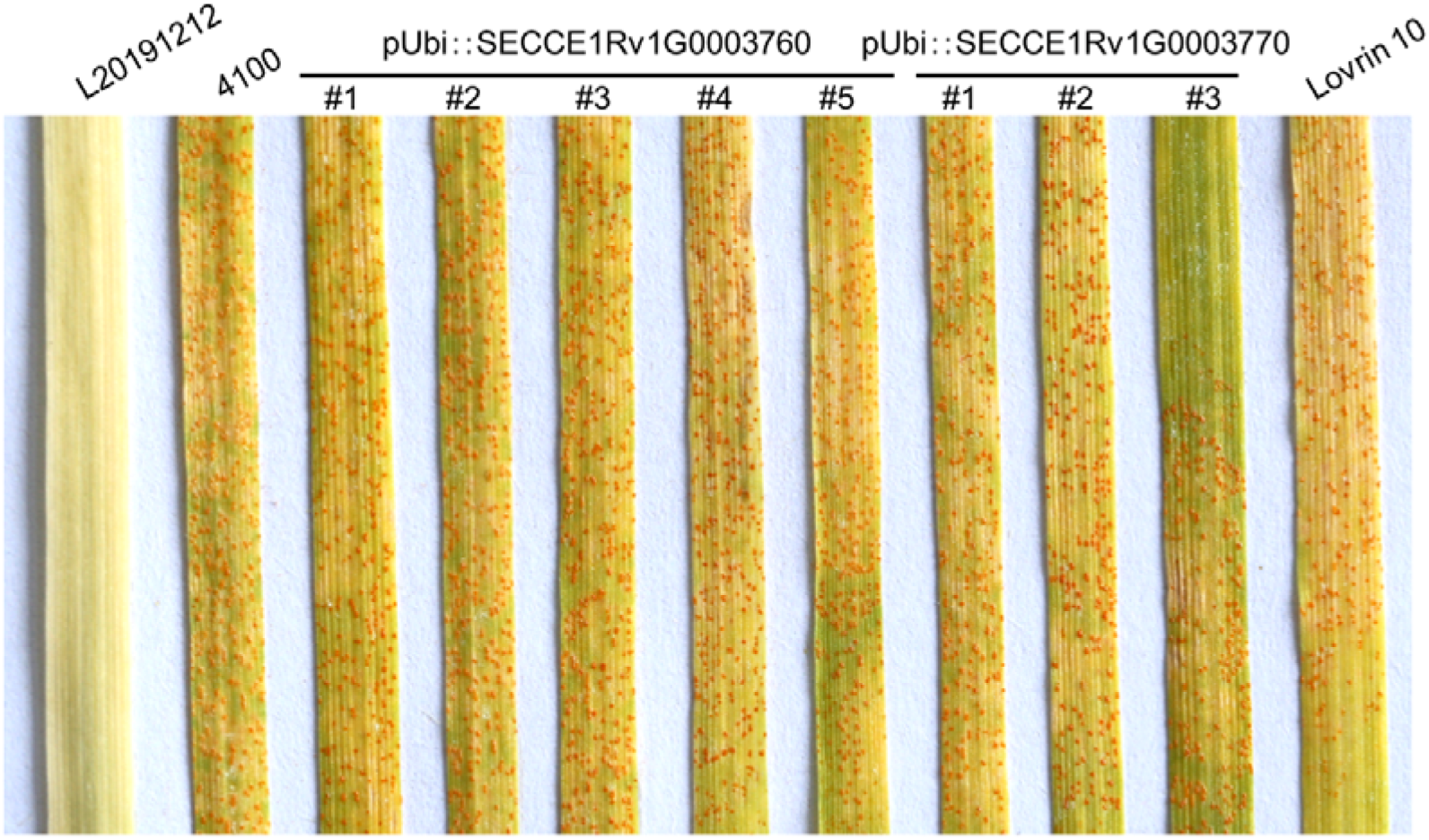
Resistance of two candidate genes to mixed physiological races of stripe rust in the field. Stripe rust mixed physiological races were collected in the field in Gansu Province, China in 2023.

**Figure S5.**
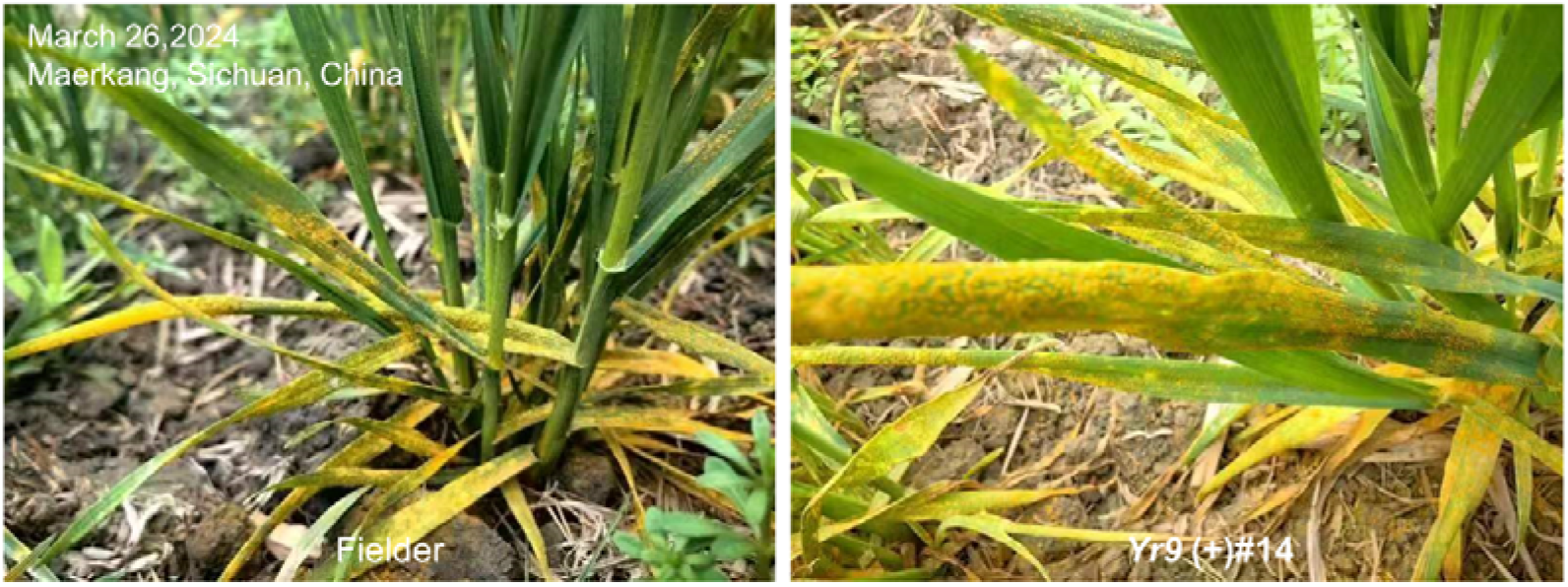
*Yr9* has insufficient stripe rust resistance under field conditions. Field phenotype of *Yr9* transgenic plants in Maerkang, Sichuan, China, compared with the common wheat Fielder.

**Figure S6.**
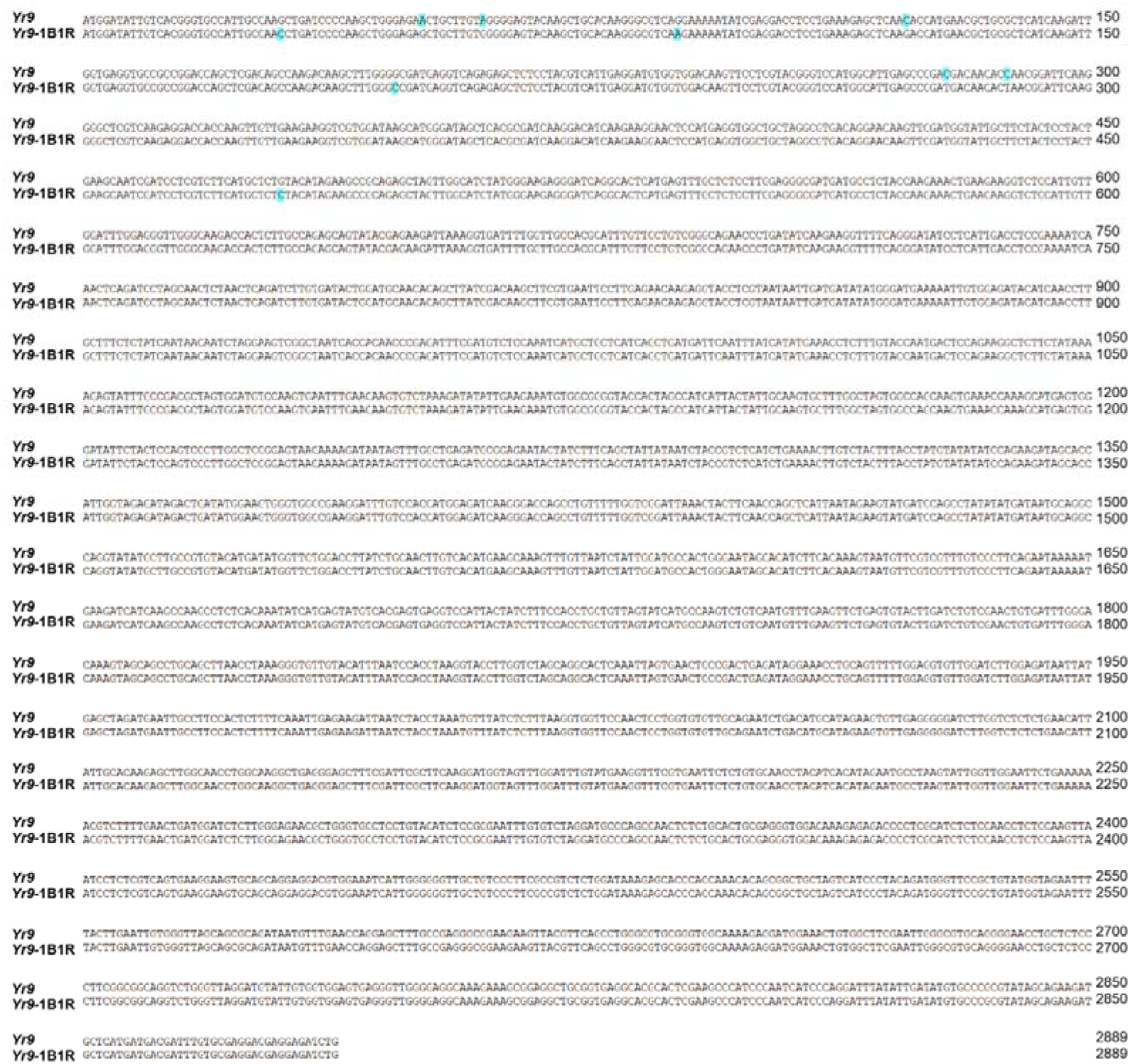
The CDS sequences of *Yr9* and *Yr9*-1B1R. A blue background highlights variable nucleotides.

**Figure S7.**
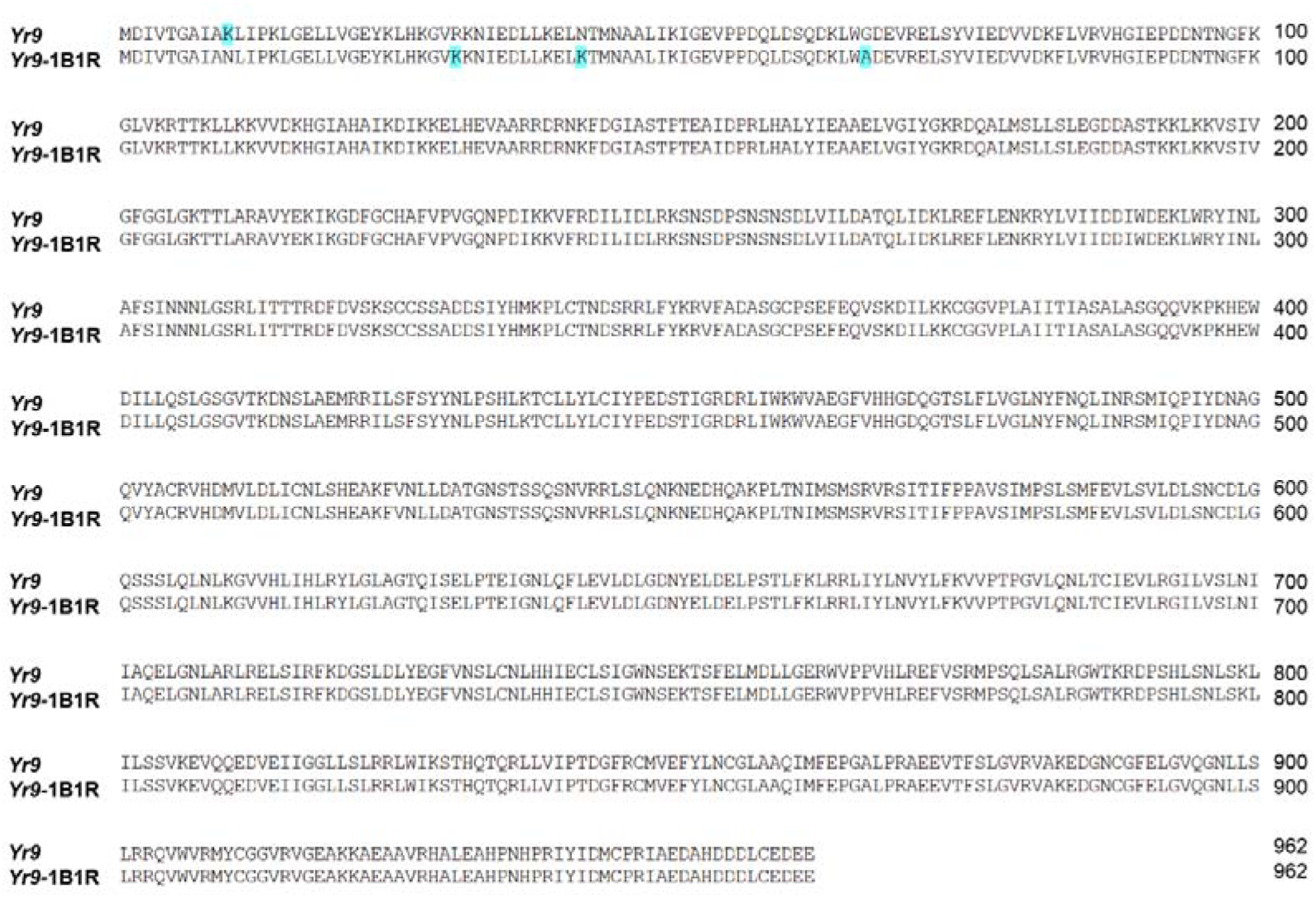
Sequence alignments of *Yr9* protein in different triticales and 1BL/1RS translocation cultivars. This figure shows the protein sequence (962 aa) alignment of *Yr9* in 21 different triticales and 1BL/1RS translocation cultivars.

### Legends for Datasets S1 to S4

Supplemental Dataset 1. InDel markers used in genetic mapping.

Supplemental Dataset 2. Transcriptome analysis of all genes in the *Yr9* interval.

Supplemental Dataset 3. Details of the triticales and 1BL/1RS translocation lines used in this paper.

Supplemental Dataset 4. Primers information used in this study.

## References

1. Chaves MS., et al., The importance for food security of maintaining rust resistance in wheat. Food Security 5(2):157–176(2013).

2. Tadesse W., Sanchez-Garcia M., Assefa SG.., Amri A, & Baum M., Genetic Gains in Wheat Breeding and Its Role in Feeding the World. Crop Breed. Genet. Genom. 1, e190005 (2019).

3. Bouvet L., et al., The evolving battle between yellow rust and wheat: implications for global food security. Theor. Appl. Genet. 135(3):741–753(2022).

4. Schwessinger B., Fundamental wheat stripe rust research in the 21 century. New Phytol. 213(4):1625–1631(2017).

5. Brown J. & Hovmøller MS., Aerial dispersal of pathogens on the global and continental scales and its impact on plant disease. Science 297(5581):537–541(2002).

6. Beddow J., et al., Research investment implications of shifts in the global geography of wheat stripe rust. Nat. Plants 1:15132(2015).

7. Wan A.M., Chen X.M., & He Z.H., Wheat stripe rust in China. Aust. J. Agri. Res. 58(6):605–619(2007).

8. Zhou Y., et al., *Triticum* population sequencing provides insights into wheat adaptation. Nat. Genet. 52(12):1412–1422(2020).

9. Consortium TIWGS, et al., A chromosome-based draft sequence of the hexaploid bread wheat (*Triticum aestivum*) genome. Science 345(6194):1251788(2014).

10. Feldman M. & Levy AA., Origin and Evolution of Wheat and Related Triticeae Species. Alien Introgression in Wheat: Cytogenetics, Molecular Biology, and Genomics, eds Molnár-Láng M, Ceoloni C, & Doležel J (Springer International Publishing, Cham), pp 21–76(2015).

11. Liu C., et al., Establishment of a set of wheat-rye addition lines with resistance to stem rust. Theor. Appl. Genet. 135(7):2469–2480(2022).

12. Wang J., et al., Centromere structure and function analysis in wheat-rye translocation lines. Plant J. 91(2):199–207(2017).

13. Huang Y., et al., Wide hybridizations reveal the robustness of functional centromeres in *Triticum-Aegilops* species complex lines. J. Genet. Genomics (2023). 10.1016/j.jgg.2023.12.001.

14. Guo X., et al., Systemic development of wheat-*Thinopyrum elongatum* translocation lines and their deployment in wheat breeding for Fusarium head blight resistance. Plant J. 114(6):1475–1489(2023).

15. Guo X.., et al., *Thinopyrum intermedium* Development and cytological characterization of wheat-translocation lines with novel stripe rust resistance gene. Front. Plant Sci. 14:1135321(2023).

16. Mater Y, et al., Linkage mapping of powdery mildew and greenbug resistance genes on recombinant 1RS from ‘Amigo’ and ‘Kavkaz’ wheat–rye translocations of chromosome 1RS.1AL. Genome 47(2):292–298(2004).

17. Mago R., et al., High-resolution mapping and mutation analysis separate the rust resistance genes *Sr31*, *Lr26* and *Yr9* on the short arm of rye chromosome 1. Theor. Appl. Genet. 112(1):41–50(2005).

18. Howell T., et al., Mapping a region within the 1RS.1BL translocation in common wheat affecting grain yield and canopy water status. Theor. Appl. Genet. 127(12):2695–2709(2014).

19. Rabinovich SV., Importance of wheat-rye translocations for breeding modern cultivar of *Triticum aestivum L*. Euphytica 100(1):323–340(1998).

20. Yang Z., et al., Utilization of 1BL/1RS translocation in wheat breeding in China. Acta Agron. Sin. 30(6):531–535(2004).

21. Yang, Z.J., and Ren, Z.L., Expression of gene *Pm8* for resistance to powdery mildew in wheat from Sichuan. J. Sichuan Agric. Univ. 15: 452–456. (1997).

22. Wang J., et al., Pan-genome analysis reveals a highly plastic genome and extensive secreted protein polymorphism in *Puccinia striiformis* f. sp. *tritici*. J. Genet. Genomics (2023).

23. Nemeth C., et al., Generation of amphidiploids from hybrids of wheat and related species from the genera *Aegilops*, *Secale*, *Thinopyrum*, and *Triticum* as a source of genetic variation for wheat improvement. Genome 58(2):71–79(2015).

24. Li Z., Li B., Tong Y.P., The contribution of distant hybridization with decaploid *Agropyron elongatum* to wheat improvement in China. J. Genet. Genomics 35(8):451–456(2008).

25. Hurni S., et al., The powdery mildew resistance gene *Pm8* derived from rye is suppressed by its wheat ortholog *Pm3*. Plant J. 79, 904–913 (2014).

26. R. Mago, et al., High-resolution mapping and mutation analysis separate the rust resistance genes *Sr31*, *Lr26* and *Yr9* on the short arm of rye chromosome 1. Theor. Appl. Genet. 112(2005).

27. Li H., et al., Cloning of the wheat leaf rust resistance gene *Lr47* introgressed from *Aegilops speltoides*. Nat. Commun. 14(1):6072(2023).

28. Ni F., et al., Sequencing trait-associated mutations to clone wheat rust-resistance gene *YrNAM*. Nat. Commun. 14(1):4353(2023).

29. Marchal C., et al., BED-domain-containing immune receptors confer diverse resistance spectra to yellow rust. Nat. Plants 4(9):662–668(2018).

30. Wen A.., et al., Genetic mapping of a major gene in triticale conferring resistance to bacterial leaf streak. Theor. Appl. Genet. 131(3):649–658(2018).

31. Alheit K, et al., Multiple-line cross QTL mapping for biomass yield and plant height in triticale (× *Triticosecale* Wittmack). Theor. Appl. Genet. 127(1):251–260(2014).

32. Dyda M., Tyrka M., Gołębiowska G., Rapacz M., & Wędzony M., Genetic mapping of adult-plant resistance genes to powdery mildew in triticale. J. Appl. Genet. 63(1):73–86(2022).

33. Shi Z.X., Chen X.M., Line.R.F., Development of resistance gene analog polymorphism markers for the *Yr9* gene resistance to wheat stripe rust. Genome 44(4):509–516(2001).

34. Chen X.M., Challenges and solutions for stripe rust control in the United States. Aust. J. Agri. Res. 58(6):648–655(2007).

35. Chen X., et al., Wheat Stripe Rust Epidemics and Races of *Puccinia striiformis* f. sp*. tritici* in the United States in 2000. Plant Dis. 86(1):39–46(2002).

36. Singh H., et al., Residual effect of defeated stripe rust resistance genes/QTLs in bread wheat against prevalent pathotypes of *Puccinia striiformis* f. sp*. tritici*. PLoS One 17(4):e0266482(2022).

37. Jin H. et al., Genetic analysis of chromosomal loci affecting the content of insoluble glutenin in common wheat. J. Genet. Genomics 42(2015).

38. Q.T. Jiang, et al., Characterization of ω-secalin genes from rye, triticale, and a wheat 1BL/1RS translocation line. J. Appl. Genet. 51(2010).

39. Kato A., Lamb JC., & Birchler JA., Chromosome painting using repetitive DNA sequences as probes for somatic chromosome identification in maize. Proc. Natl. Acad. Sci. U.S.A. 101(37):13554–13559(2004).

40 Huang Y., et al., New insights on the evolution of nucleolar dominance in newly resynthesized hexaploid wheat *Triticum zhukovskyi*. Plant J. 115(5):1298–1315(2023).

41. Rabanus-Wallace M., et al., Chromosome-scale genome assembly provides insights into rye biology, evolution and agronomic potential. Nat. Genet. 53(4):564–573(2021).

42. Maccaferri M., et al., Durum wheat genome highlights past domestication signatures and future improvement targets. Nat. Genet. 51(5):885–895(2019).

43. A.M. Bolger, M. Lohse, B. Usadel T., Trimmomatic: a flexible trimmer for Illumina sequence data. Bioinformatics 30(15):2114–2120(2014).

44. Dobin A., et al., STAR: ultrafast universal RNA-seq aligner. Bioinformatics 29(1):15–21(2013).

45. McKenna A., et al., The Genome Analysis Toolkit: a MapReduce framework for analyzing next-generation DNA sequencing data. Genome Res. 20(9):1297–1303(2010).

46. Mansfeld B. & Grumet R., QTLseqr: An R Package for Bulk Segregant Analysis with Next-Generation Sequencing. Plant Genome 11(2)(2018).

47. Pertea M., Kim D., Pertea G., Leek J., & Salzberg SL., Transcript-level expression analysis of RNA-seq experiments with HISAT, StringTie and Ballgown. Nat. Protoc. 11(9):1650–1667(2016).

48. Kim D., Paggi JM., Park C., Bennett C., & Salzberg SL., Graph-based genome alignment and genotyping with HISAT2 and HISAT-genotype. Nat. Biotechnol. 37(8):907–915(2019).

49. Pertea M., et al., StringTie enables improved reconstruction of a transcriptome from RNA-seq reads. Nat. Biotechnol. 33(3):290–295(2015).

50. Frazee AC., et al., Ballgown bridges the gap between transcriptome assembly and expression analysis. Nat. Biotechnol. 33(3):243–246(2015).

51. Guo W., et al., Origin and adaptation to high altitude of Tibetan semi-wild wheat. Nat. Commun. 11(1):5085(2020).

52. Li H. & Durbin R., Fast and accurate short read alignment with Burrows-Wheeler transform. Bioinformatics 25(14):1754–1760(2009).

53. Meng L., Li H., Zhang L., & Wang J., QTL IciMapping: Integrated software for genetic linkage map construction and quantitative trait locus mapping in biparental populations. Crop J. 3(3):269–283(2015).

54. Bolser D., Staines D., Perry E., & Kersey PJ., Ensembl Plants: Integrating Tools for Visualizing, Mining, and Analyzing Plant Genomic Data. Methods Mol. Biol. 1533:1–31(2017).

